# Analysis of meiotic recombination in *Drosophila simulans* shows heterozygous inversions do not cause an interchromosomal effect

**DOI:** 10.1101/2024.03.09.584235

**Authors:** Bowen Man, Elizabeth Kim, Alekhya Vadlakonda, David L. Stern, K. Nicole Crown

## Abstract

Chromosome inversions are of unique importance in the evolution of genomes and species because when heterozygous with a standard arrangement chromosome, they suppress meiotic crossovers within the inversion. In Drosophila species, heterozygous inversions also cause the interchromosomal effect, whereby the presence of a heterozygous inversion induces a dramatic increase in crossover frequencies in the remainder of the genome within a single meiosis. To date, the interchromosomal effect has been studied exclusively in species that also have high frequencies of inversions in wild populations. We took advantage of a recently developed approach for generating inversions in *Drosophila simulans*, a species that does not have inversions in wild populations, to ask if there is an interchromosomal effect. We used the existing chromosome 3R balancer and generated a new chromosome 2L balancer to assay for the interchromosomal effect genetically and cytologically. We found no evidence of an interchromosomal effect in *D. simulans*. To gain insight into the underlying mechanistic reasons, we qualitatively analyzed the relationship between meiotic double-strand break formation and synaptonemal complex assembly. We find that the synaptonemal complex is assembled prior to double-strand break formation as in *D. melanogaster*; however, we show that the synaptonemal complex is assembled prior to localization of the oocyte determination factor Orb, whereas in *D. melanogaster*, synaptonemal complex formation does not begin until Orb is localized. Together, our data show heterozygous inversions in *D. simulans* do not induce an interchromosomal effect and that there are differences in the developmental programming of the early stages of meiosis.

## Intro

Chromosome inversions are of unique importance in the evolution of genomes and species because when heterozygous with a standard arrangement chromosome in an individual, they effectively suppress meiotic crossover formation within the inversion (1–3). This crossover suppression happens two ways: first, there is a slight decrease in the frequency of crossover recombination within the interval; second, crossovers within an inversion result in chromosome rearrangements that cause aneuploid (and thus inviable) gametes, preventing inheritance of the recombinant chromosome in the next generation (1–3). Crossover suppression has consequences for fertility on the single generation time scale because crossovers are essential for meiotic chromosome segregation and over long evolutionary timescales because suppressing crossovers affects the rate of gene flow within and across populations (4, 5).

In Drosophila species, heterozygous inversions have an additional effect during meiosis termed the interchromosomal effect, whereby the presence of a heterozygous inversion induces a dramatic increase in crossover frequencies in the remainder of the genome in a single generation (comprehensively reviewed in (6)). In *Drosophila melanogaster*, the interchromosomal effect is caused by activating the pachytene checkpoint and extending the developmental window where prophase (and crossover formation) is occurring, ultimately leading to a 50-500% increase in crossover frequencies on the non-inverted chromosomes (7). The increase in crossover frequencies is not caused by increasing the number of meiotic DNA double-stranded breaks (DSBs); rather there is a shift in the balance between crossover and noncrossover repair outcome during meiotic recombination (8).

The molecular consequences of heterozygous inversions and the interchromosomal effect have far ranging impacts on the rate of gene flow within and between populations: gene flow is reduced locally between the inversion and the standard arrangement chromosome but is increased in the remainder of the genome. Since many Drosophila species are polymorphic for inversions in wild populations, understanding the molecular mechanisms and the pervasiveness of the interchromosomal effect is important for understanding how inversions shape populations. Various estimates have shown that inversions occur at very high frequencies in multiple Drosophila species. Perhaps the most informative estimate is the number of species within a subgenus that are monomorphic or polymorphic for inversions (9, 10). Within the Drosophila subgenus, approximately half of species are polymorphic for inversions (41 species) and half are monomorphic (44 species). Within the Sophophora subgenus, 38 species are polymorphic for inversions (including *D. melanogaster*), while only 5 are monomorphic (*D. simulans*, *D. mauritiana*, *D. sechellia*, *D. erecta*, and *D. santomea*). It is striking that within the *D. melanogaster* species subgroup, the simulans species complex is monomorphic (*D. simulans*, *D. mauritiana*, and *D. sechellia*), while their sister species *D. melanogaster* is highly polymorphic (9, 10).

It is surprising that such closely related species within the *D. melanogaster* species subgroup have dramatically different frequencies of inversions. Previous work has attempted to explain this difference at the molecular level by asking if *D. simulans* and *D. melanogaster* differ in their response to repairing DSBs and found conflicting results. Woodruff and Ashburner irradiated males with 4000 rads of ionizing radiation and found the same number of chromosome aberrations (translocations and inversions) in both species (11). However, Lemke performed the same experiment and found chromosome aberrations in 24.8% of offspring from *D. melanogaster* and only 0.8% from *D. simulans* (12). A later experiment by Inoue also found about half as many chromosome breaks in *D. simulans* compared to *D. melanogaster* after exposure to ionizing radiation (13). Additionally, *D. simulans* has been resistant to karyotype manipulation in the lab. Despite exhaustive attempts to generate balancer chromosomes using the same methods that have been successful in *D. melanogaster*, all attempts that we are aware of have failed (14).

Recently, a doubly inverted chromosome 3R balancer (*j3RM1*) was successfully generated in *D. simulans* using an elaborate site-specific recombination approach that screened for loss and gain of fluorescent markers, which allowed 5,000-10,000 flies to be screened in each step of the process (14). In the work presented here, we generated an additional chromosome 2L doubly inverted balancer (*j2LM1*) and use it and *j3RM1* to ask if the interchromosomal effect is present in a species that is normally monomorphic for inversions in the wild. Using recessive marker scoring to determine crossover frequencies, we found no evidence of an interchromosomal effect. We also found no evidence of pachytene checkpoint activation in response to heterozygous inversions in this species, additionally suggesting there is no interchromosomal effect. We do find differences in the developmental timing of early meiosis in *D. simulans* compared to *D. melanogaster* and suggest that this difference is one of many possible explanations for why there is not an interchromosomal effect in this species.

## Methods

### Drosophila stocks and husbandry

All *D. simulans* stocks were maintained at 25 degrees Celsius on standard cornmeal media. *w^501^* (stock #14021-0251.11), *y^1^ v^2^ f^1^ bb^1^* (stock #14021-0251.147), and *jv^1^ st^1^ e^1^ p^1^* (stock #14021-0251.174) stocks were obtained from the National Drosophila Species Stock Center. We note that chromosome 3R in *D. melanogaster* is inverted relative to *D. simulans* (15), so the gene order for *e* and *p* is reversed relative to *D. melanogaster*. In our hands, *bb^1^* was no longer visible in the *y^1^ v^2^ f^1^ bb^1^* stock. Kenya C157.4 was a gift from Amanda Larracuente. The third chromosome inversion *j3RM1* was previously published and is maintained as *dsx/j3RM1* (14).

### Generating the *j2LM1* balancer stock

We followed the protocol described in Stern (2022) to generate two overlapping inversions on chromosome arm 2L in *D. simulans*. Four plasmids carrying reporter genes expressing variously colored fluorescent proteins in different anatomical regions and either FRT or KD yeast recombination sites were integrated separately into existing attP landing sites in *D. simulans* (16): sim-2810 p{ie1-dsRed::FRT-RC,attB} into sim-952 p{attP, 3XP3-EYFP} inserted at 2L: 830,996; sim-2720 p{MHC-EGFP::KD} into sim-1048 p{attP, 3XP3-EYFP} inserted at 2L: 6,392,252; sim-2664 p{3XP3-DsRed::FRT} into sim-1230 p{attP, 3XP3-EYFP} inserted at 2L: 13,194,453; and sim-2719 p{MHC-DsRed::KD-RC} into sim-960 p{attP, 3XP3-EYFP} inserted at 2L: 22,086,768. All genome coordinates are relative to *D. simulans* genome assembly NCBI:GCA_016746395.1. All four transgenes were recombined onto a single chromosome arm. Flies carrying all four transgenes on a single chromosome were crossed to flies carrying a source of heat-shock inducible Flipase p(MUH-HES-FLPL, w+, attB) integrated into 1029-simD-234.5 at 3R: 17,461,328. Plastic vials containing developing larvae in their food media were placed in a 37°C bead bath (Lab Armor) for 1 hour, followed by 1 hour at room temperature, and a second 37°C heat shock for one hour. Male progeny were crossed to a *w-* strain of *D. simulans* and offspring of this cross were screened for loss of ie1-DsRed and 3XP3-dsRed, but retention of MHC-EGFP and MHC-DsRed, which indicates inversion between the FRT sites. Flies carrying this singly-inverted chromosome were then crossed to flies carrying a source of heat-shock inducible KD recombinase p(MUH-HES-KD::PEST, w+, attB) integrated into 2176-simD-299.6 at 2L:6,583,842. This attP landing site is derived from 1048-simD-299.6, but has had its 3XP3-EYFP marker inactivated by CRISPR-Cas9 mutagenesis of the EYFP gene (16). Larvae from this cross were heat shocked as described above, and adults were crossed to a *w-* strain of *D. simulans*. Offspring were screened for loss of MHC-EGFP and MHC-DsRed, but presence of 3XP3-EYFP, which is present in all four landing sites with integrated recombination plasmids, but not in the landing site with the KD-recombinase integrated. These progeny carried presumptive double-inversions on chromosome 2L, which we named *j2LM1*. To determine whether *j2LM1* might serve as a useful balancer chromosome, flies carrying this chromosome, which could be recognized by the 3XP3-EYFP markers, were backcrossed to a *w-* strain of *D. mauritiana* for 10 generations. Twelve independent backcrosses were performed and DNA was prepared from one individual of the tenth generation from each backcross and subjected to multiplexed-shotgun genotyping to estimate chromosome ancestry (17). All progeny were heterozygous for chromosome 2L, indicating that *j2LM1* prevented inheritance of recombination events on chromosome 2L (Figure 1).

**Figure 1.**
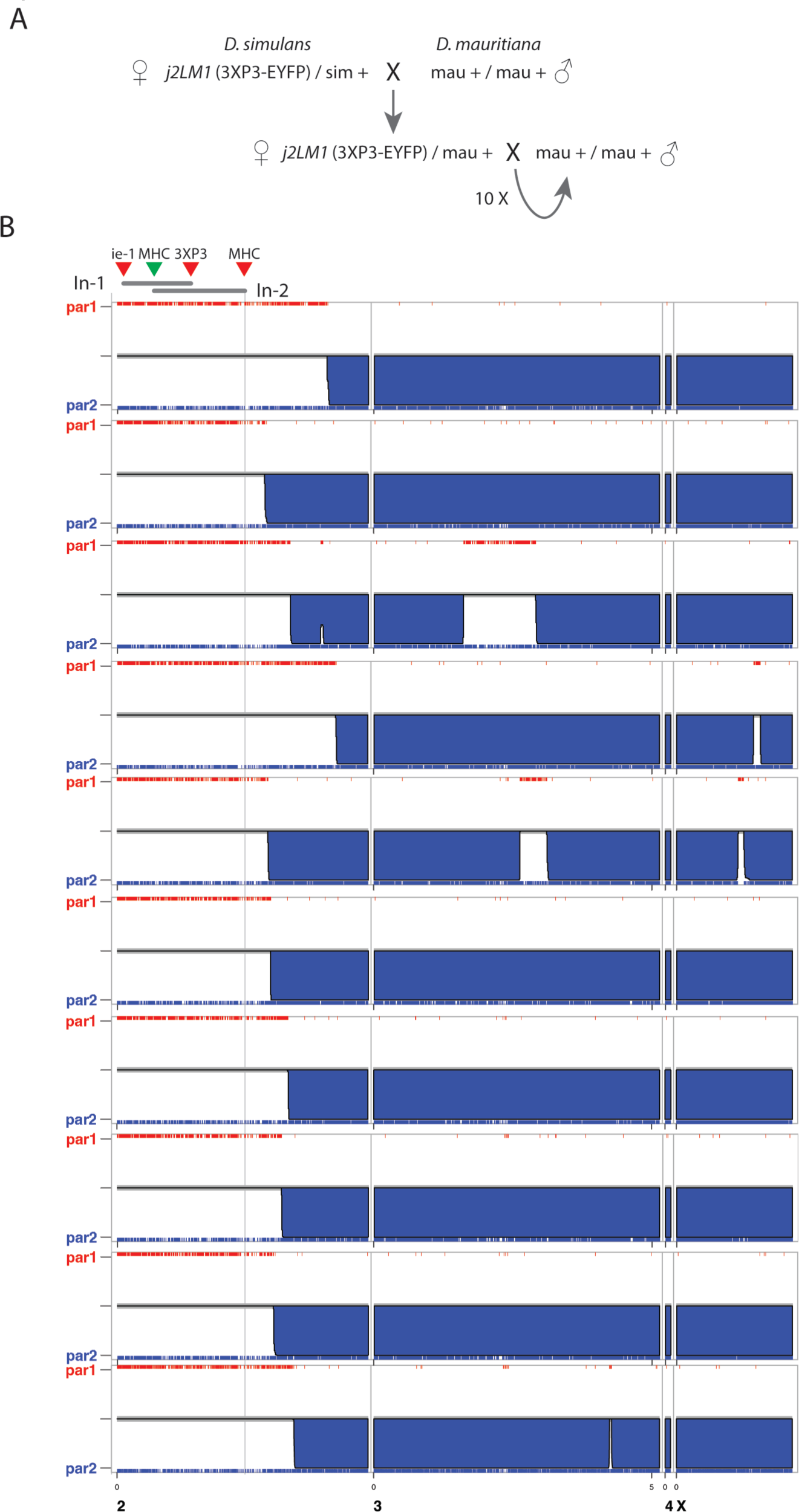
The *D. simulans j2LM1* balancer chromosome suppresses meiotic crossovers. A) *D. simulans j2LM1* females were crossed to *D. mauritiana* males. The F1 offspring were then backcrossed to *D. mauritiana* for 10 generations. B) Chromosomal ancestry estimates for individual flies from 10 different backcrossed lines. The y-axis represents the probability that a region is homozygous for *D. simulans* (red) or *D. mauritiana* (blue) SNPs. Regions that appear blank on the y-axis represent regions that are heterozygous for *D. simulans* and *D. mauritiana* SNPS. Individual SNP markers are shown on the top and bottom x-axes. Chromosome positions and scale bar are listed at the bottom. The *j2LM1* inversion breakpoints are shown above as colored triangles, and the entire span of the inversions is shown as gray horizontal bars.

### Immunofluoresence

2-3 day old mated females were put on yeast paste overnight. Ovaries were dissected in PBS and fixed for 20 minutes in 1000 μl of solution containing 2% paraformaldehyde (Ted Pella cat. no. 18505), 0.5% Nonidet P-40 (Sigma cat. no. I8896), 200 μl PBS and 600 μl heptane. Ovaries were then washed three times for ten minutes each in PBS with 0.1% Tween-20 (PBST), blocked for one hour at room temperature in PBS with 1% BSA (MP Biomedicals cat. no. 152401) and incubated with primary antibody diluted in PBST overnight at 4 degrees. Ovaries were then washed three times in PBST and incubated in secondary antibody diluted in PBST for 4 hours at room temperature. DAPI was added for the last 10 minutes at a concentration of 1 μl/ml. Ovaries were washed again three times for 15 minutes each in PBST. All wash steps and antibody incubations were done while nutating. Ovaries were mounted in ProLong Glass (Invitrogen cat. no. P36980) and allowed to cure for the manufacturer’s suggested time.

Since *D. simulans* is closely related to *D. melanogaster*, we reasoned that some antibodies against *D. melanogaster* proteins would work in *D. simulans*. We found that the following *D. melanogaster* antibodies result in similar staining patterns in *D. simulans* as in *D. melanogaster*: anti-Corolla (18), anti-phospho-H2AV (19), and anti-Orb (20). We used rabbit anti-Corolla at 1:3000 (gift from Scott Hawley), mouse anti-phospho-H2AV at 1:500 (DSHB clone UNC93-5.2.1), and mouse anti-Orb at 1:20 (DSHB clone 6H4 at 1:40 combined with clone 4H8 at 1:40). We used the following secondary antibodies: Goat anti-Mouse IgG2b Alexa Fluor 594 (Thermofisher # A-21145), Goat anti-Mouse IgG1 Alexa Fluor 488 (Thermofisher #A-21121), and Goat anti-Rabbit Alexa Fluor 647 (Thermofisher #A-21244).

Ovaries were imaged on a Leica Stellaris 5 confocal microscope using an HC PL APO 63x/1.4 NA Oil objective. Images were acquired using the Lighting module with an Airy pinhole size of 0.75 AU and the standard default settings dictated by the pinhole size. All images were deconvolved using the Leica Lightning internal software with default settings.

### Crossover scoring

We used recessive marker scoring to analyze crossover frequencies. We crossed *jv st e p/+* females to *jv st e p* homozygous males and scored offspring for each of the recessive markers. Crossovers were identified as locations where the genotype switched from mutant to wildtype. These crosses were performed in a wildtype background or in females that were heterozygous for a balancer. Some stocks contained a *white* mutation, and in these cases, only female offspring were scored. The same crosses were repeated to measure crossover frequencies in the X chromosome using a *y v f bb* chromosome.

The *j2LM1* stock is maintained over a wildtype chromosome 2, which allowed us to control for genetic background by measuring crossover frequencies in wildtype and inversion siblings from the same cross. We crossed *w-*; +/*j2LM1* females to *y v f b* or *jv st e p* males. We recovered females that were heterozygous for the marker chromosome and either heterozygous for *j2LM1* (marked by 3xP3-GFP) or homozygous for the standard arrangement chromosome (marked by the absence of 3xP3-GFP). We crossed these females to *y v f b* or *jv st e p* males and scored crossovers in the offspring.

### Statistics

Crossover frequencies were compared between two genotypes using Fisher’s exact test and Bonferonni’s correction for multiple tests. When three genotypes were compared simultaneously, a Chi-square test was performed. All statistical tests were performed in GraphPad Prism version 10.2 for Mac (GraphPad Software, Boston, MA USA).

## Results

### Generating inversions in *D. simulans*

To ask if there is an interchromosomal effect in *D. simulans,* we took advantage of a multiply inverted third chromosome balancer that was recently generated (14). This balancer, termed *j3RM1*, has two nested inversions that cover the entirety of chromosome 3R (Figure 2). Importantly, the left arm of this chromosome is not inverted and thus it only balances the right arm. We also generated a new chromosome 2L balancer chromosome (*j2LM1*) that has two nested inversions on the left arm of chromosome 2 (see Methods for details). Similarly to *j3RM1*, the right arm of *j2LM1* is not inverted (Figure 2).

**Figure 2.**
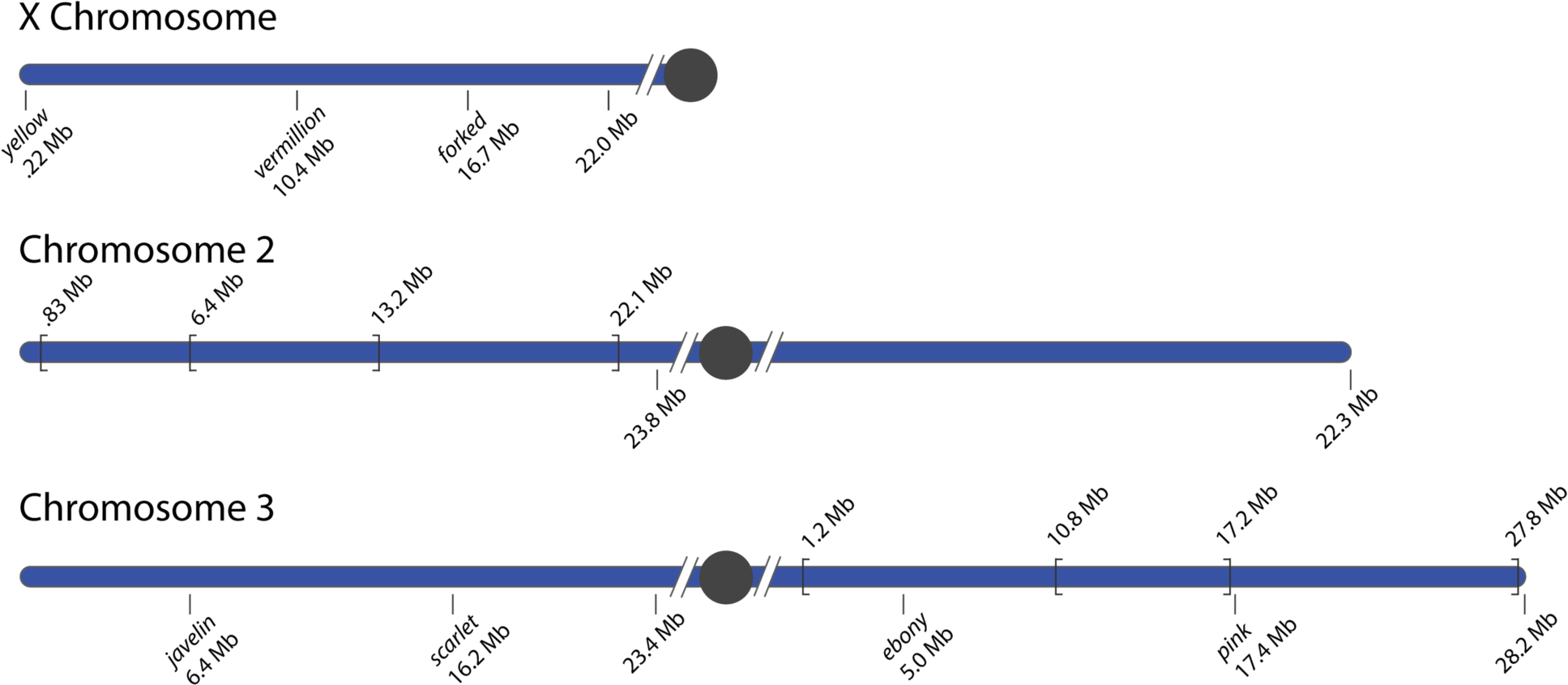
Chromosome sizes, positions of the recessive markers, and location of inversion breakpoints used in this study. Recessive markers and their physical position are shown below the chromosomes. Inversion breakpoints are depicted as brackets and their physical position is listed above the chromosome. The end of each chromosome assembly is shown below the chromosome. All physical positions were obtained from the *D. simulans* genome NCBI:GCF_016746395.2 Black circles represent the location of centromere. Hashed lines represent unassembled pericentric heterochromatin of unknown size.

To ensure that *j2LM1* does indeed suppress the inheritance of crossing-over events, we crossed females heterozygous for *j2LM1* to its sister species *D. mauritiana*, and then backcrossed the F1 offspring to *D. mauritiana* for ten generations in 12 independent lines. Thus, recombination between *D. simulans and D. mauritiana* chromosomes was possible in every generation. We sequenced one fly from each of the 12 lines and determined if they were heterozygous or homozygous for *D. simulans* and *D. mauritiana* single nucleotide polymorphisms (SNPs). Regions where heterozygosity was maintained for ten generations indicated crossovers did not form or were not inherited. We found each of the 12 lines was heterozygous for *D. simulans* and *D. mauritiana* SNPS in the portion of chromosome 2L with the inversion (Figure 1). Most of the remainder of the genome in each individual was homozygous for *D. mauritiana* SNPs.

Although we previously showed that *j3RM1* suppresses crossovers using the same *D. mauritiana* backcross scheme just described, we used recessive marker scoring to confirm that this balancer suppresses crossovers within species. We crossed *j3RM1* to the multiply marked scoring chromosome containing *jv st e p* and found no crossovers in the *e – p* interval, which is included within the *j3RM1* inverted regions (Table 1 and Supplemental Table 1). Crossover frequencies were statistically the same as wildtype in the *jv – st* and *st – e* intervals, showing that there is no intrachromosomal effect from inversions as has been occasionally reported for *D. melanogaster* (2, 21, 22).

**Table 1.**
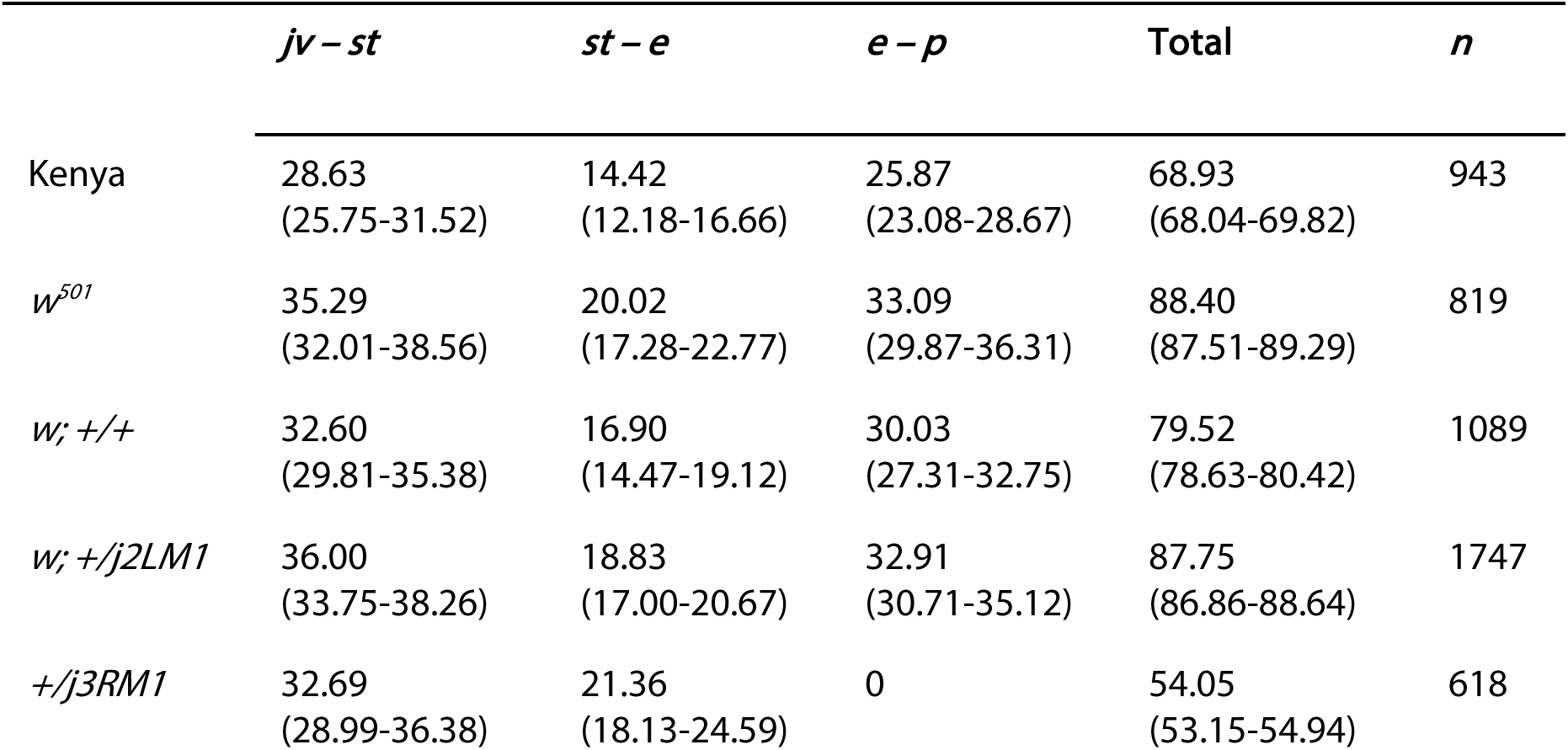
Crossover frequencies in cM for each interval along chromosome 3 in varying genotypes. *w*; +/+ is the wildtype full sibling to *w*; +/*j2LM1*. Values below cM are 95% confidence intervals. The variations in map length between Kenya, *w^501^*, and the wildtype full sibling strain are statistically significantly different (Chi-square, p < 0.0001).

### Significant variation in crossover rates between wildtype stocks

*D. melanogaster* shows considerable intraspecies variation in crossover rates (23). To determine crossover rates in various wildtype *D. simulans* strains, we used recessive marker scoring on chromosome 3 in three different wildtype strains: Kenya C167.4, *w^501^*, and a full sibling wildtype strain derived from the *j2LM1* stock. We found statistically significant variation in crossover frequencies across all three genotypes (Table 1 and Supplemental Table 1, Chi-square test, p < 0.0001), with an overall map length of 68.93 cM in Kenya C167.4, 88.40 cM in *w^501^* and 79.52 cM in the full sibling wildtype strain. The variation in crossover frequencies occurs in all three individual intervals and no one interval can account for all variation. This variation in crossover frequencies between wildtype strains, combined with the recent discovery of high levels of pericentromeric genome instability in *w^501^* (24), led us to compare crossover frequencies in wildtype full siblings whenever possible.

### There is no interchromosomal effect in *D. simulans*

To determine if there is an interchromosomal effect, we measured crossover frequencies on chromosome 3 in females heterozygous for the *j2LM1* inversion and in wildtype full siblings derived from the same cross. Based on the *D. melanogaster* historical literature, we *a priori* defined the interchromosomal effect as at least a 50% increase in crossover frequency in individual intervals or total map length (6). We found that crossover frequencies on chromosome 3 are not statistically significantly different in individual intervals, but the entire map length is statistically longer (Table 1 and Supplemental Table 1. All p-values reported in Supplemental Table 3). However, the increase in total map length is only 110% of the wildtype map length, thus there is no interchromosomal effect.

Because the regulation of recombination can be different between autosomes and sex chromosomes (25), we next asked if there is an interchromosomal effect on the X chromosome. We measured crossover frequencies on the X chromosome in *j3RM1* heterozygotes, *j2LM1* heterozygotes, and wildtype full siblings to *j2LM1.* We did not compare crossover frequencies to a wildtype full sibling of *j3RM1* because this stock is maintained over a *doublesex* mutation. Like chromosome 3, the crossover frequencies in individual intervals were not statistically different between inversion heterozygotes and the wildtype full sibling nor was the entire map length (Table 2 and Supplemental Table 2. All p-values reported in Supplemental Table 3). Together, the data from chromosome 3 and the X chromosome suggest there is not an interchromosomal effect in *D. simulans*.

**Table 2.**
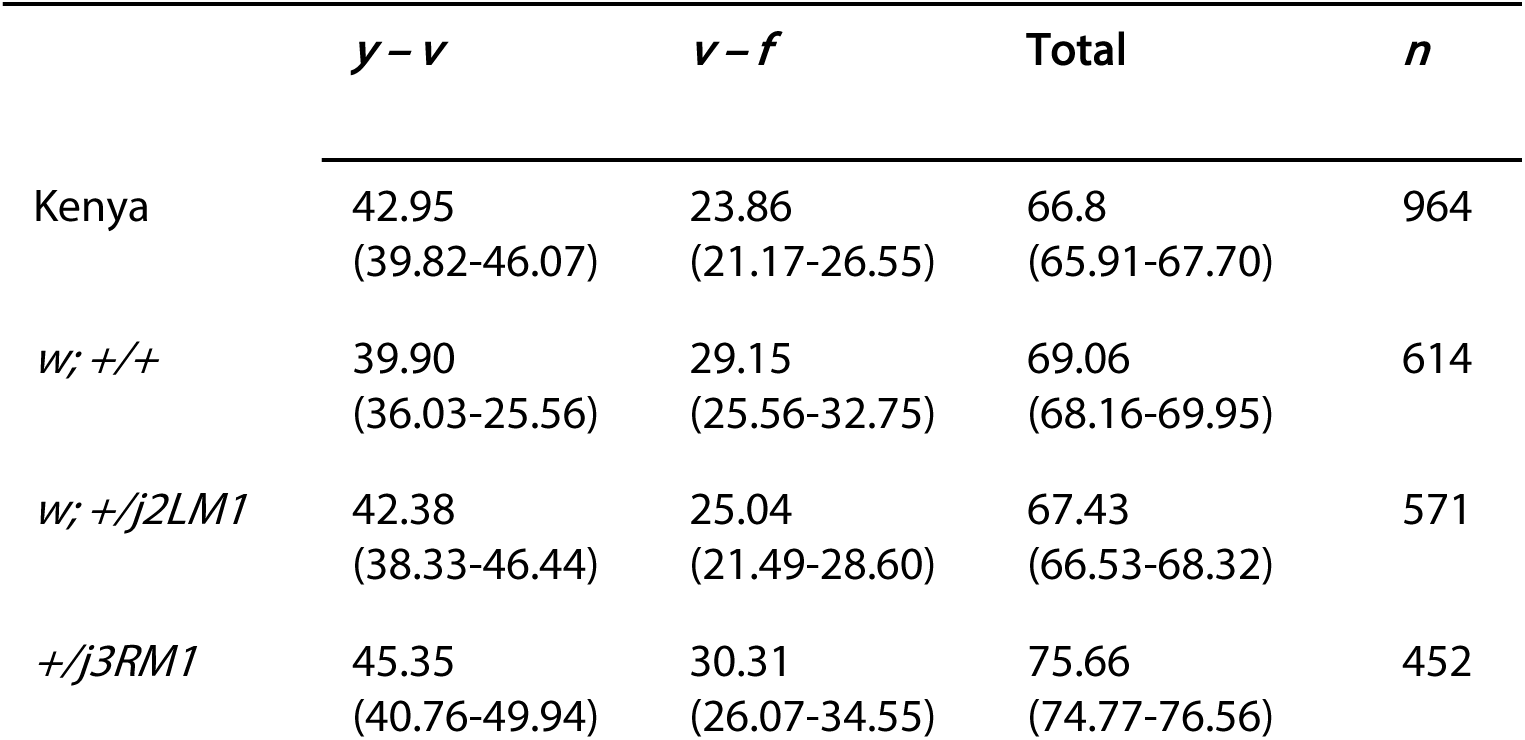
Crossover frequencies in cM for each interval along the X chromosome in varying genotypes. *w*; +/+ is the wildtype full sibling to *w*; +/*j2LM1*. Values below cM are 95% confidence intervals. No intervals are statistically significantly different (Fisher’s exact test with Bonferonni correction for multiple tests). All p-values are shown in Supplemental Table 3.

### Heterozygous inversions do not cause a delay in oocyte specification

In *D. melanogaster*, meiosis occurs within the context of a 16-cell cyst that progresses through the germarium. Up to four nuclei within each cyst initiate meiosis by building synaptonemal complex in region 2a, but by region 2b only two nuclei remain in meiosis (the pro-oocytes), and by the end of the germarium in region 3, one of the pro-oocytes exits meiosis and the remaining cell is specified as the oocyte (26). The RNA-binding protein Orb is essential for oocyte specification; this is reflected in the protein localization of Orb as it is initially present in the cytoplasm of all cells within a cyst beginning in region 2a but becomes concentrated in the oocyte during specification (20). During the interchromosomal effect in *D. melanogaster,* heterozygous inversions activate the pachytene checkpoint, which monitors recombination intermediates and chromosome axes (7, 27). Activating the pachytene checkpoint causes a prolonged prophase and a delay in oocyte specification, resulting in two Orb-positive oocytes instead of one in region 3 (7).

To ask if there is a 2-oocyte phenotype in inversion heterozygotes in *D. simulans,* we stained germarium with the *D. melanogaster* anti-Orb antibody (20). While we do not have *orb* mutants in *D. simulans* to test for antibody specificity, the Orb staining pattern is identical to *D. melanogaster* (Figure 3a). Like in *D. melanogaster,* Orb localizes to the cytoplasm of all cysts starting in region 2a, begins to concentrate in two cells starting in region 2b, and is concentrated in one cell only in region 3. In both the wildtype full sibling and *j2LM1* heterozygotes, only 3% of region 3 germaria had two oocytes (Figure 3b). This again suggests that there is no interchromosomal effect in *D. simulans*.

**Figure 3.**
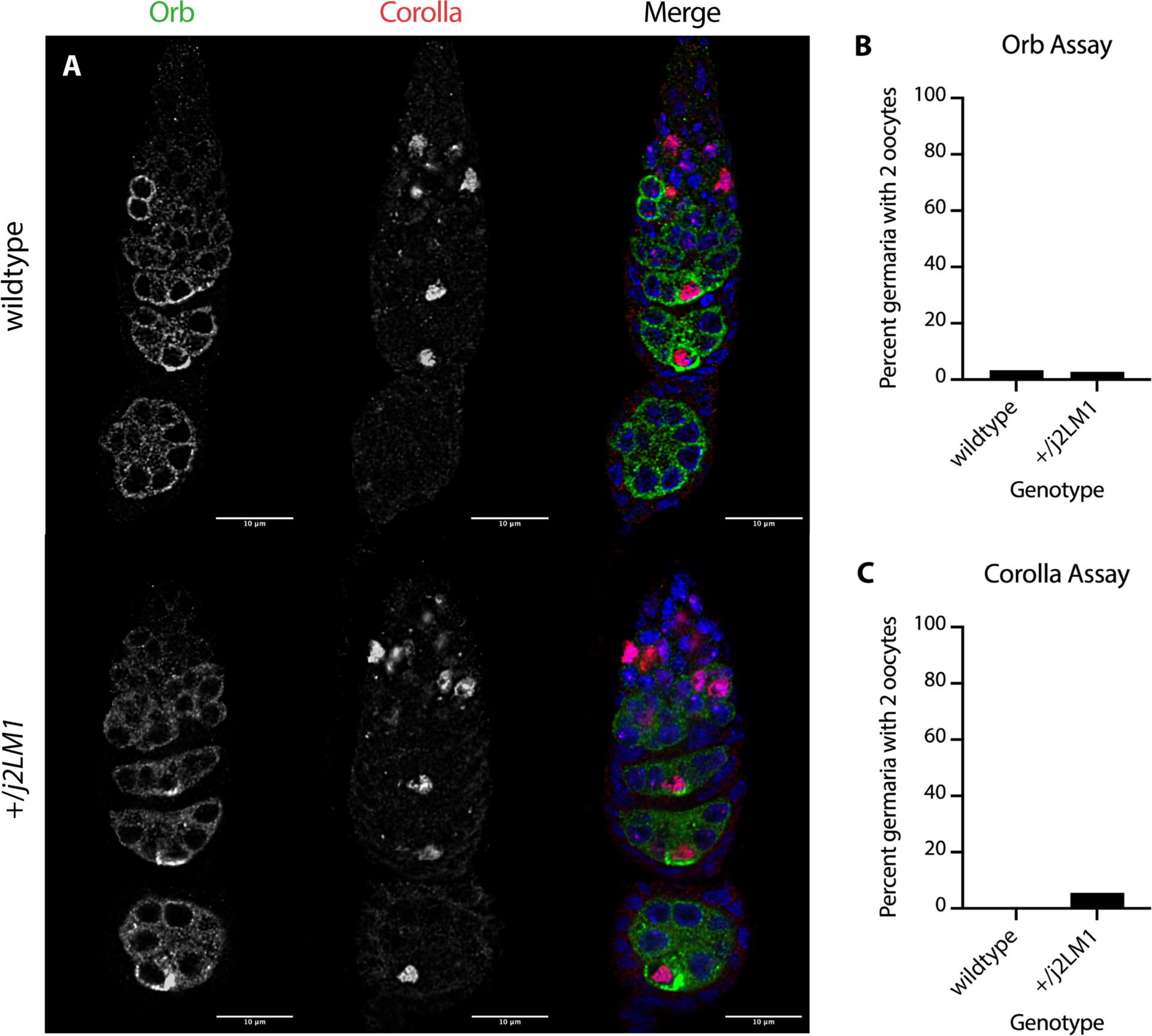
Analysis of Orb and Corolla localization in full sibling wildtype and *j2LM1* heterozygotes shows no delay in oocyte specification. A) Immunofluorescence in *D. simulans* germaria and stage 2 egg chambers with antibodies against Orb and Corolla in full sibling wildtype (top) and +/*j2LM1* heterozygotes (bottom). Orb accumulates in identical patterns to *D. melanogaster*. It first appears in the cytoplasm of all cells of a 16-cell cyst in region 2a. By region 2b, it has started to concentrate in the pro-oocytes and by region 3 is concentrated in the designated oocyte. Different from *D. melanogaster*, Corolla is loaded onto full length synaptonemal complex at the end of region 1, before Orb signal is detected. Note that because a single z-slice shown, some signal is out of the focal plane. B and C) Quantification of the 2-oocyte phenotype using Orb localization (B) and Corolla localization (C). There are no statistically significant differences in the frequency of two oocytes in region 3 (Fisher’s exact test, p = 0.15)

During oocyte specification, one of the two pro-oocytes exits meiosis and disassembles the synaptonemal complex (26). In *D. melanogaster,* the 2-oocyte phenotype can also be measured by the number of nuclei that have full length synaptonemal complex present in region 3. When measured this way, approximately 10% of wildtype germaria have two synaptonemal complex-positive nuclei, but 50-90% of germaria in inversion heterozygotes have two synaptonemal complex-positive nuclei in region 3 (7). Since assaying for two oocytes using the synaptonemal complex as a marker is more sensitive than using Orb, we stained *D. simulans* germaria with the *D. melanogaster* antibody against the synaptonemal complex component Corolla (Figure 3a). Although we do not have a *corolla* mutant to test for antibody specificity, we found that the nuclear staining pattern is identical to that in *D. melanogaster* and clearly identifies the synaptonemal complex in this species (Figure 3a). We also found that, like *D. melanogaster,* multiple cells within a cyst initiate meiosis and build synaptonemal complex, but by region 3, 100% of germaria had one nucleus with full-length synaptonemal complex. In the *j2LM1* heterozygotes, we saw the same pattern of Corolla staining as in wildtype. There was a marginal increase in the number of germaria with two Corolla-positive nuclei (Figure 3c), but this increase was not statistically significant (Fisher’s exact test, p = 0.15). Together, the similar crossover frequencies between wildtype and inversion heterozygotes and the lack of a delay in oocyte specification show that there is no interchromosomal effect in *D. simulans*.

### Relationship between double-strand break formation and the synaptonemal complex in *D. simulans*

To understand the molecular reasons why there is not an interchromosomal effect in *D. simulans*, we qualitatively analyzed the relationship between DNA double-strand break (DSB) formation and the synaptonemal complex. In mouse and *S. cerevisiae,* synaptonemal complex formation is genetically dependent on DSB formation, while in *D. melanogaster* and *C. elegans,* synaptonemal complex formation is genetically independent of DSBs (28–33). The relative relationship between the synaptonemal complex and DSB formation can result in very different physical environments where recombination is occurring. In *D. melanogaster*, the timing of these events is such that the synaptonemal complex is fully assembled before DSBs form and thus recombination takes place within the context of a fully assembled synaptonemal complex (34). We reasoned that since the synaptonemal complex is thought to be important for implementing crossover patterning mechanisms, and that crossover patterning mechanisms are partially responsible for the interchromosomal effect (8), the relationship between DSBs and the synaptonemal complex might be different in *D. simulans* than in *D. melanogaster*.

We were able to qualitatively analyze the relative timing of DSB formation and synaptonemal formation in *D. simulans* through immunostaining. To visualize the synaptonemal complex, we stained with the *D. melanogaster* antibody against Corolla, a central region protein (18), and found staining consistent with that in *D. melanogaster* (Figures 3 and 4). DSBs can be visualized cytologically using antibodies against phosphorylated H2AV (19, 34). We tested the mouse monoclonal *D. melanogaster* antibody against phospho-H2AV (19) in *D. simulans* and, despite a moderate level of background staining, found a nuclear localization pattern consistent with that of meiotic DSBs in *D. melanogaster* (Figure 4).

**Figure 4.**
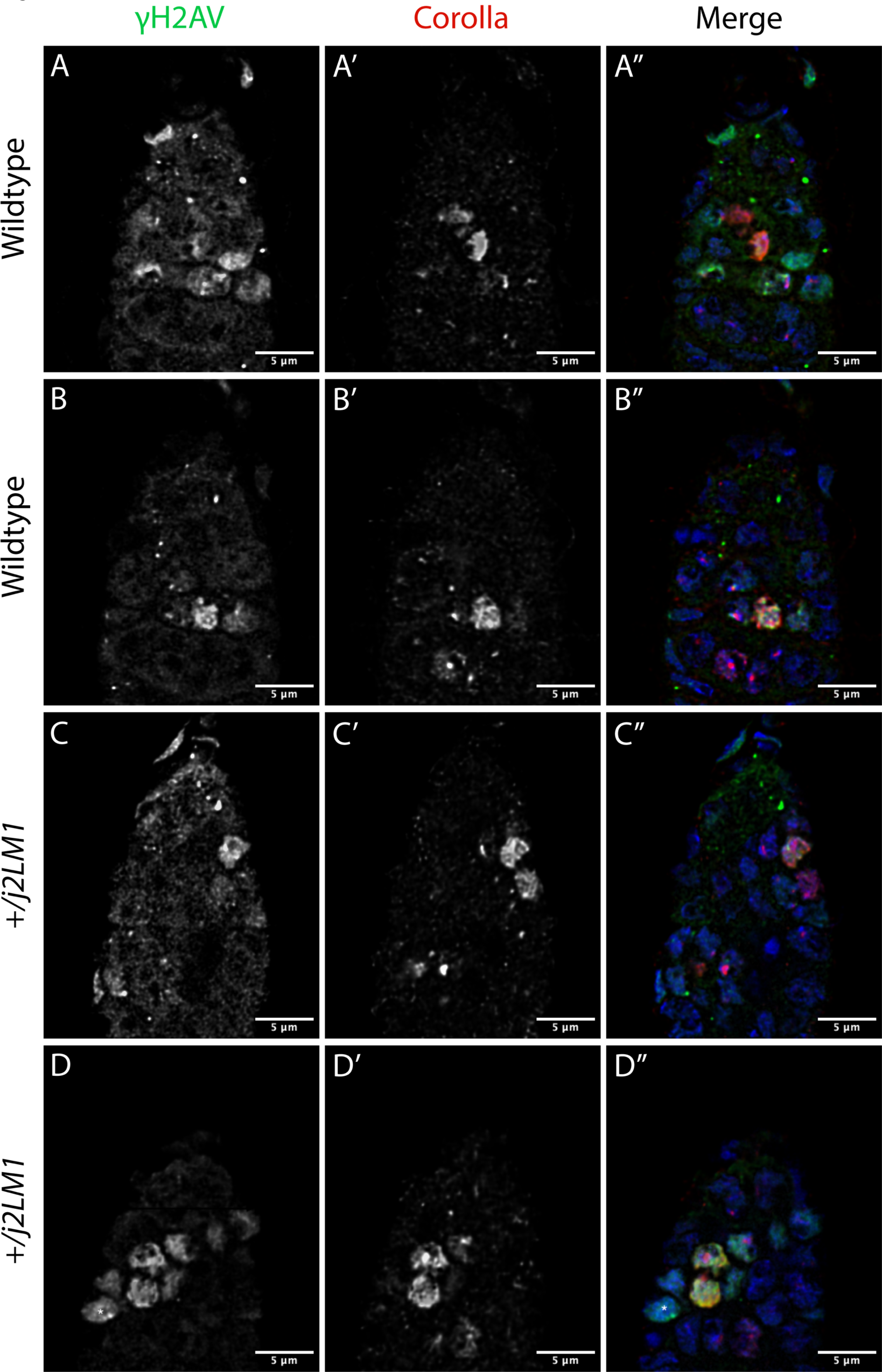
Qualitative analysis of DSB and synaptonemal complex formation suggests the synaptonemal complex forms prior to DSB formation in *D. simulans*. A) A single z-slice in the early germarium of a wildtype full sibling showing the earliest pair of Corolla-positive nuclei. One of the nuclei has faint phospho-H2AV signal while the other nucleus shows none. B) A different single z-slice of the same germarium in A, showing that the next cyst in that germarium shows robust phospho-H2AV signal. C) A single z-slice in the early germarium of a *j2LM1* heterozygote showing the same pattern as wildtype: the earliest pair of Corolla-positive nuclei shows phospho-H2AV signal in one of the nuclei but not the other. D) A different single z-slice of the same germarium in C, showing the next cyst has robust phospho-H2AV signal. The asterisk denotes a nurse cell that has phospho-H2AV signal without Corolla signal.

In *D. melanogaster*, the synaptonemal complex begins assembling in early region 2A at the same time as Orb appears, and full-length synaptonemal complex is present by late region 2a (35–37). Interestingly, in *D. simulans*, we found full length Corolla staining at the end of region 1 before Orb appears (Figure 3). It is possible that these nuclei are at the very end of zygotene and the cysts are on the cusp of entering region 2a, but regardless of if they are late zygotene or early pachytene, the developmental timing of SC assembly in *D. simulans* represents a marked deviation from the timing in *D. melanogaster*. Additionally, the Corolla-positive nuclei always appear in pairs, suggesting that two cells per cyst enter meiosis (Figure 4). Like *D. melanogaster*, full length Corolla staining is visible in two nuclei in region 2b and in one nucleus in region 3 (Figure 3).

In *D. melanogaster,* DSBs are initiated in region 2a quickly after the synaptonemal complex is fully assembled (19, 26, 34). To determine the relative timing of DSB and synaptonemal complex formation in *D. simulans,* we analyzed phospho-H2AV patterns in the early germarium (Figure 4). We observed that in the earliest pairs of nuclei with full length Corolla, one of the nuclei had faint phospho-H2AV foci while the other nucleus had no phospho-H2AV staining (Figure 4a and 4c). By region 2a, all Corolla positive nuclei had robust phospho-H2AV staining (Figure 4b and 4d). We did detect nuclei with phospho-H2AV but without Corolla (Figure 4c and 4d, asterisks); these are most likely the nurse cells as it is known that nurse cells undergo DSB formation in *D. melanogaster* (34). We repeated these same qualitative analyses in the *j2LM1* heterozgyotes and found no differences in the localization patterns of Corolla or phospho-H2AV (Figure 4c and 4d). We interpret these staining patterns such that full length synaptonemal complex is built prior to robust DSB formation in *D. simulans*. Thus, while the relative timing of DSB and SC formation is similar between *D. melanogaster* and *D. simulans*, the early stages of prophase occur prior to Orb localization in *D. simulans*. Determining whether synaptonemal complex assembly is genetically dependent on DSB formation in *D. simulans* will require future genetic analyses.

## Discussion

We set out to determine if there is an interchromosomal effect in *D. simulans* because the lack of inversions in wild and lab populations suggest it may be an interesting comparative model for understanding how heterozygous inversions shape gene flow dynamics and how, at the molecular level, mechanisms of recombination and crossover regulation might differ. We found no evidence supporting the presence of an interchromosomal effect in *D. simulans*, either genetically or cytologically. One limitation of our approach is that only one arm of chromosomes 2 or 3 were inverted. It is possible that crossovers were able to form on the chromosome arms that contained the standard arrangement and that a crossover on one arm of a chromosome prevents an interchromosomal effect. We do not think this is the case based on data from *D. melanogaster* showing a single inversion on chromosome 2L was able to cause an interchromosomal effect (38). Another limitation of our study is the large genetic intervals we used for determining crossover frequencies. There are a limited number of visible phenotypic markers available in *D. simulans* and, for the current study, we were limited to what is available. While the genetic intervals were large, they spanned regions of the chromosomes where the interchromosomal effect is the strongest in *D. melanogaster* – the pericentromeric and subtelomeric regions (6). In particular, the marker distribution on chromosome 3 would have been sufficient to detect increases in crossover frequencies across the centromere. The marker distribution on the X chromosome prevented us from detecting crossovers near the centromere, but the interchromosomal effect is strong at the distal end of the X chromosome in *D. melanogaster* and our markers did span that portion of the chromosome. Thus, the marker distribution was sufficient to detect an interchromosomal effect.

It is unclear what the function of the interchromosomal effect is. Is it an evolved response in species with high numbers of segregating inversions or is it just an indirect result of checkpoint monitoring of normal meiotic processes? Looking at the interchromosomal effect in other Drosophila species can provide some insight. For example, *D. persimilis* and *D. pseudoobscura* are two hybridizing species with inversions between them; in interspecies crosses where the female hybrids are heterozygous for an inversion, there is a strong interchromosomal effect with characteristics similar to that in *D. melanogaster* (39). Additionally, *D. robusta*, which has segregating inversions in natural populations, also displays an interchromosomal effect (6, 40). The work presented here is the first analysis of the interchromosomal effect in a monomorphic species and we have shown that there is not an interchromosomal effect, perhaps suggesting that the interchromosomal effect is not present in species where it is not needed. Future work analyzing the interchromosomal effect in other monomorphic species, such as *D. mauritiana* or *D. sechellia*, should give insight into this question (41). Recent advances in high-efficiency CRISPR approaches make these experiments possible (42).

One possible mechanistic reason there is no interchromosomal effect in *D. simulans* is that crossover rates are higher in this species compared to *D. melanogaster* and there may simply not be much room for increasing crossover rates. True et al (43) directly compared crossover rates in *D. melanogaster* and *D. simulans* using recessive marker scoring and found that the total genetic map length in *D. simulans* is 1.3 times longer than in *D. melanogaster*. More specifically, the total map length of the X chromosome is identical between species, but map lengths of chromosomes 2 and 3 are significantly longer. A more recent analysis used whole genome sequencing to determine crossover rates in wild caught *D. melanogaster* and *D. simulans* trios (44). They found that overall crossover rates were 2.06 cM/Mb in a European population of *D. melanogaster*, 3.44 cM/Mb in a West African population of *D. melanogaster*, and 3.04 cM/Mb in a European *D. simulans* population.

Related to the increased rates of recombination is the observation that the centromere effect is weaker in *D. simulans*. In most species, crossovers are suppressed near centromeres – a phenomenon called the centromere effect (45). Interestingly, True et al showed that the centromere effect is weaker in *D. simulans* compared to *D. melanogaster,* and in fact, the increase in crossover frequencies in *D. simulans* can be mostly attributed to higher crossover frequencies across the centromere, as opposed to on the chromosome arms (43). Wang et al were able to precisely map the location of crossovers using a sequencing approach and found that in *D. melanogaster*, the closest crossover to a centromere occurred 6.8 Mb away (of 177 crossovers examined), but in *D. simulans* the closest crossover was 1.3 Mb from a centromere (of 109 crossovers examined) (44). Together, these data suggest that the centromere effect is weaker in *D. simulans* than in *D. melanogaster*. The interchromosomal effect has the strongest impact on crossover frequencies in the centromere spanning intervals (6), thus it could be possible that if there is a weaker centromere effect in *D. simulans* – that is, crossovers are already higher in these regions – there simply is no room to increase crossover rates.

Assessing the interchromosomal effect cytologically allows us to rule out higher crossover rates as the reason there is not an interchromosomal effect. The interchromosomal effect in *D. melanogaster* is caused by activating the pachytene checkpoint, which then allows more crossovers to form (7). If heterozygous inversions activate the pachytene checkpoint in *D. simulans*, but more crossovers do not form because crossover rates are already at or near a maximum level, we still would have detected the 2-ooctye phenotype. Thus, we do not think higher crossover rates and a weaker centromere effect are the reason there is no interchromosomal effect in *D. simulans*.

We asked if there are differences in the timing of DSB and synaptonemal complex formation that might explain why there is no interchromosomal effect in *D. simulans*. At the resolution of immunostaining, we found that the synaptonemal complex forms before DSBs as it does in *D. melanogaster*. However, there is a clear difference in the developmental timing of zygotene in *D. simulans* because the synaptonemal complex is fully assembled before Orb localization. The pachytene checkpoint delays oocyte specification and extends the window during which crossovers can form, ultimately leading to higher crossover rates during the interchromosomal effect (7). The correlation between oocyte specification and meiotic chromosome dynamics raises the possibility that crossover control mechanisms are active during certain developmental windows that are related to, or can be marked by, oocyte specification. If the initial steps of crossover and synaptonemal complex formation are occurring prior to that developmental window in *D. simulans*, then perhaps this prevents checkpoint activation and an interchromosomal effect. This hypothesis remains to be tested and will require an in-depth characterization of meiosis in *D. simulans*.

## Data availability

All stocks are available upon request. The authors affirm that all data necessary for confirming the conclusions of the article are present within the article, figures, and tables.

## Acknowledgements

We thank Scott Hawley for antibodies, Amanda Larracuente for stocks, and the National Drosophila Species Stock Center for stocks.

---

**Supplemental Table 1.**
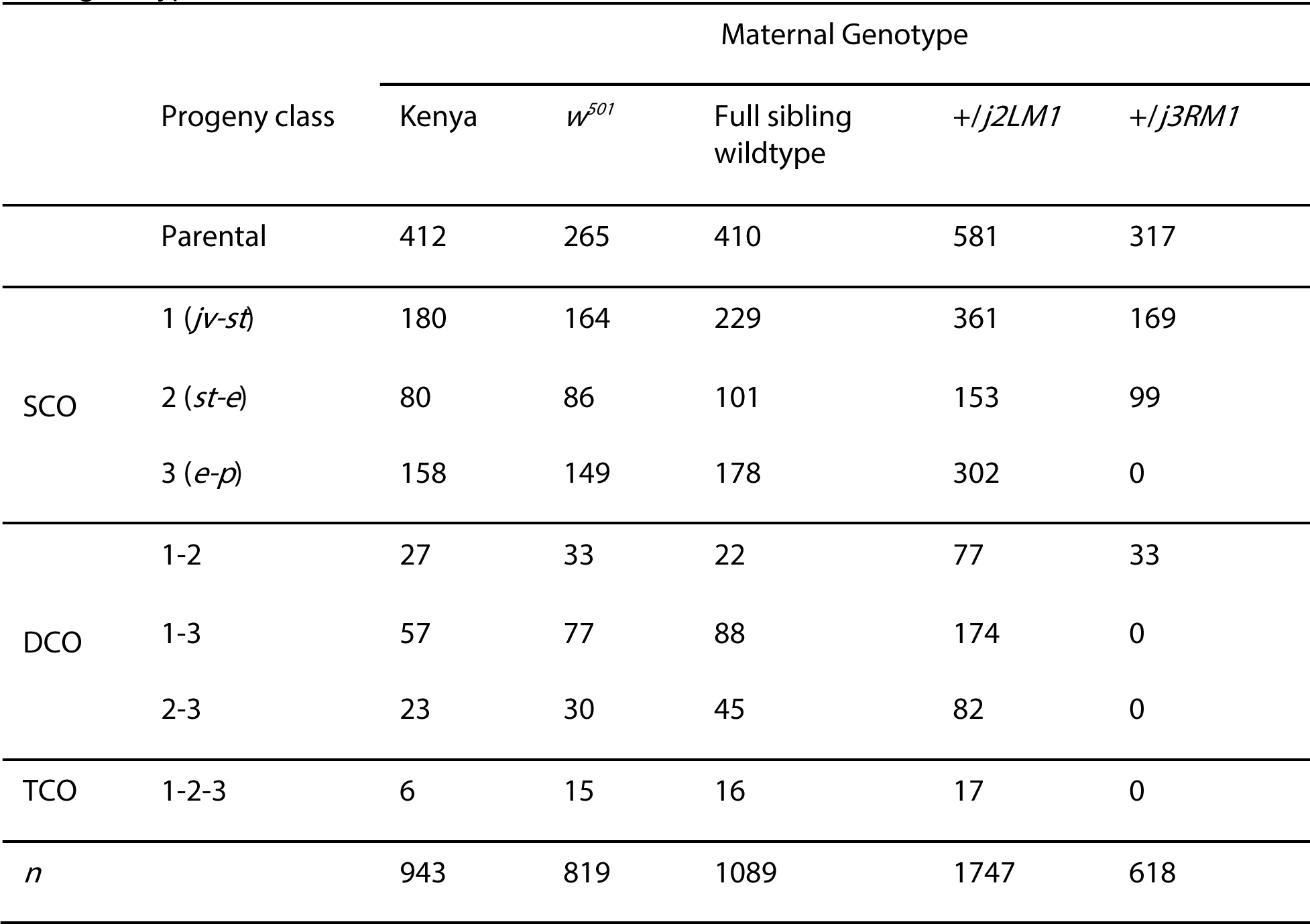
Raw numbers of meiotic crossovers on chromosome 3. Shown are the number of single crossovers (SC), double crossovers (DCO), and triple crossovers (TCOs) for each genotype in each interval. Total number of flies counted is listed in the bottom row for each genotype.

**Supplemental Table 2.** Raw numbers of meiotic crossovers on the X chromosome. Shown are the number of single crossovers (SCOs) and double crossovers (DCOs) for each genotype in each interval. Total number of flies counted is listed in the bottom row for each genotype.

|  |  | Maternal Genotype |  |  |  |
| --- | --- | --- | --- | --- | --- |
| Progeny class |  | Kenya | Full sibling<br>wildtype | *+/*j2LM1** | *+/*j3RM1** |
| Parental |  | 393 | 246 | 246 | 180 |
| SCO | 1 (*y-f*) | 341 | 189 | 182 | 135 |
|  | 2 (*f-v*) | 157 | 123 | 83 | 67 |
| DCO | 1-2 | 73 | 56 | 60 | 70 |
| *n* |  | 964 | 614 | 571 | 452 |

**Supplemental Table 3.**
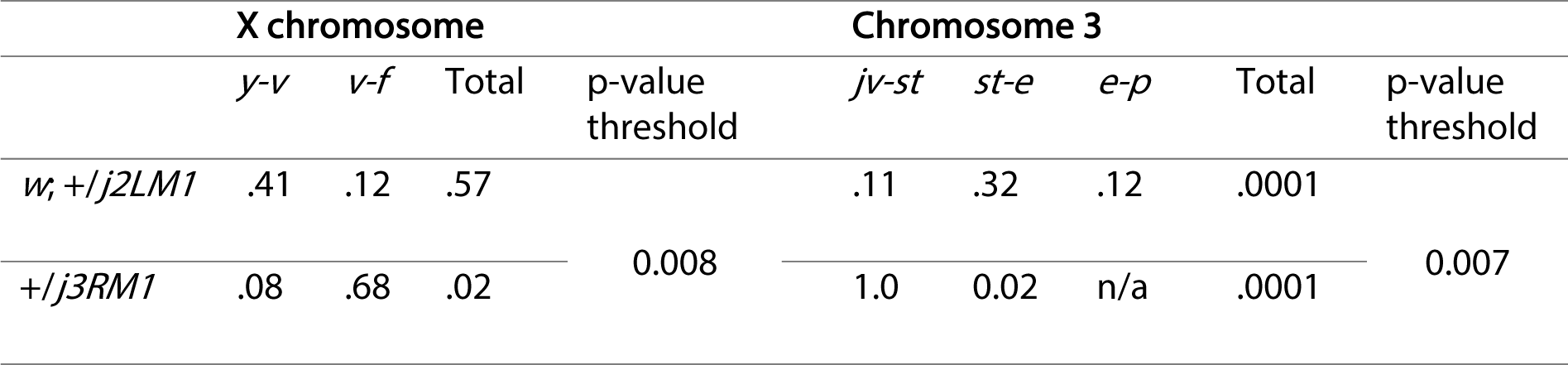
P-values for map length comparisons between full sibling wildtype of *j2LM1* and all other inversion heterozygotes using Fisher’s exact test with Bonferonni correction for multiple tests. P value thresholds for significance from Bonferonni correction are shown.

